# AT1R and Integrin β3 Synergize to Drive Aortic Dissection via Non-Canonical Wnt/β-Catenin Signaling

**DOI:** 10.1101/2025.03.26.645618

**Authors:** Hongcheng Jiang, Zixuan Liu, Yu Li, Xiaodan Zhong, Yunkun Qu, Shiliang Li, Lei Dai, Ying Zhang, Mengwen Wang, Wenjun Zhang, Wanjun Liu, Xingwei He, Wei Dong, Thati Madhusudhan, Zhiyong Li, Hongjie Wang, Hesong Zeng

## Abstract

Aortic dissection (AD), a life-threatening cardiovascular emergency, continues to impose high mortality due to insufficient therapeutic options, as monotherapy targeting angiotensin II type 1 receptor (AT1R) demonstrates limited clinical efficacy. Utilizing single-cell RNA sequencing, we identified integrin β3 as a critical driver of AD progression, with expression levels positively correlated with disease severity. Histopathological validation in human AD specimens and a murine angiotensin II (AngII)-infusion model confirmed marked upregulation of integrin β3 activation. Pharmacological blockade of integrin β3 with Cyclo(-RGDfK) significantly attenuated aortic pathogenesis *in vivo*, reducing dissection incidence and aortic degeneration. Mechanistically, AngII-mediated AT1R activation induced formation of a receptor complex with integrin β3, triggering its conformational activation. Transcriptomic profiling revealed that activated integrin β3 potentiates vascular endothelial dysfunction by binding glycogen synthase kinase 3β (GSK3β), which stabilizes β-catenin via a non-canonical Wnt signaling axis. This pathway drives endothelial barrier disruption, hallmarks of aortic wall destabilization in AD. Our findings unveil a previously unrecognized synergy between AT1R and integrin β3, implicating aberrant Wnt/β-catenin signaling as a nexus of endothelial injury in AD pathogenesis. These results advocate for a paradigm-shifting dual-therapeutic strategy concurrently targeting AT1R and integrin β3 to restore vascular homeostasis, offering a mechanistically grounded approach to mitigate this lethal disease. This work bridges critical gaps in understanding AD pathophysiology and provides a transformative framework for precision therapeutics.

## Introduction

Aortic dissection (AD) represents a critical medical emergency characterized by delamination of the aortic wall layers, culminating in catastrophic morbidity and mortality rates exceeding 50% within 48 hours if untreated [1, 2]. Despite advances in surgical interventions, pharmacological strategies to mitigate AD progression remain limited, underscoring an urgent need to elucidate its molecular drivers [3]. The aortic wall comprises of three layers, namely the intima, media, and adventitia [4], among which endothelial cells (ECs) critically regulate vascular barrier and integrity of the vascular intima [5, 6]. Pathologically, ECs are vulnerable to reactive oxidative stress (ROS), inflammatory cytokines, and aberrant shear forces, which collectively induce apoptosis, intimal damage, and eventual dissection [7, 8]. However, the molecular mechanisms linking EC dysfunction to AD pathogenesis remain incompletely defined, hampering therapeutic innovation.

Composed of 18 α- and 8 β-subunits, functionally distinct integrin heterodimers are transmembrane receptors bridging the cells and extracellular matrix (ECM) and help cells cope with various internal and external perturbations [9–11]. Among these, integrin αvβ3—a prominent RGD-binding receptor—orchestrates EC survival, angiogenesis, and matrix remodeling [12, 13]. Notably, integrin β3 has been implicated in modulating matrix metalloproteinase-9 (MMP9), a protease central to vascular wall degradation and aortic aneurysm formation [14, 15]. These findings suggest a potential role for integrin β3 dysregulation in AD progression, though its precise contributions to EC dysfunction and aortic destabilization remain unexplored.

As a member of the G protein-coupled receptor (GPCR) family, angiotensin II type 1 receptor (AT1R) is a is a well-established mediator of vascular hypertrophy, ROS, and inflammation—hallmarks of AD pathogenesis [16, 17]. However, meta-analysis of randomized controlled trials found angiotensin converting enzyme inhibitor (ACEi) and angiotensin receptor blocker (ARB) prescription alone fail to attenuate aortic dilatation in AD, implying the involvement of non-canonical AT1R signaling pathways or synergistic molecular partners [18]. Intriguingly, integrin β3 exhibits bidirectional crosstalk with multiple GPCRs, including protease-activated receptors (PARs) and growth factor receptors, via direct interactions or shared downstream effectors such as RhoA and MMPs [19–21]. Additionally, the interaction of integrin β3 cytoplasmic domain with G protein Gα13 and Gα12 suggests its involvement in mediating non-canonical G protein signaling. Furthermore, we have previously shown that integrin β3 crosstalk with PAR3 temporally activates RhoA activation in renal epithelial cells, suggesting a broader role in GPCR signaling modulation [22–24]. We thus hypothesize that integrin β3 may act as a co-receptor or amplifier of AT1R-driven pathological signaling in ECs, contributing to AD progression.

Here, we investigate the role of integrin β3 in AD pathogenesis through complementary clinical, experimental, and mechanistic approaches. First, we demonstrate upregulated integrin β3 activation in ECs from human AD specimens and murine models, correlating with MMP9 expression and intimal disruption. Using an integrin β3-selective inhibitor (Cyclo-RGDfK), we show that targeting integrin β3 attenuates AD incidence and aortic remodeling in mice.

Furthermore, co-immunoprecipitation and transcriptomic profiling reveal a physical interaction between integrin β3 and AT1R, alongside enrichment of non-canonical pathways involving Wnt/β-catenin signaling. These findings position integrin β3 as a novel modulator of AT1R signaling in AD, offering a mechanistic foundation for dual-target therapeutic strategies to counteract this lethal condition.

## Method

### RNA Sequencing and Bioinformatics Analysis

Human umbilical vein endothelial cells (HUVECs, NC versus integrin β3 knockdown) were treated with angiotensin II (AngII, 1 μM, 24 h) or vehicle control (n = 3/group). Total RNA was extracted by an RNA purification kit (Magen, China). The concentration and quality of RNA were determined using a NanoDrop spectrophotometer (KAIAO, China). Sequencing service were performed by Wefindbio Biotechnology Co., Ltd. (Wuhan, China). Limma (R package, version 3.56.2) was used to identify differentially expressed genes (DEGs). GSEA software (version 4.3.2) was applied to probe the significant functional differences between different groups. The gene sets “m5.go.v2023.2.Mm.symbols.gmt” was obtained from the GSEA website MsigDB database (http://www.gsea-msigdb.orggsea/msigdb/). Gene sets were considered as significantly enriched by the adjusted p-value < 0.05 and |normalized enrichment score (NES)| > 1. Data analysis was performed by R software (version 4.1.3).

### Single cell RNA Sequencing analysis

Public single-cell RNA-seq dataset GSE222318 (4 acute AD, 3 subacute AD, 2 chronic AD, 5 controls) was reanalyzed using *Seurat* (v4.0) [25, 26]. Cells were filtered based on the criteria: nFeature RNA > 600, nFeature RNA < 6000, and percent.mt < 10. The FindClusters function was applied with resolution = 0.5 to identify distinct cell populations. Cell clustering was conducted based on previous classification criteria [25], and the AddModuleScore function was used to compute pathway scores for different cell types.

### Animal experiments

The animal procedures conform to the guidelines from Directive 2010/63/EU of the European Parliament on the protection of animal used for scientific purpose and all animal studies were conducted in accordance with the standard procedures approved by the Institutional Animal Care and Use Committee of Tongji Hospital, Huazhong University of Science and Technology (Issue Number: 4154). Male 6-week-old ApoE^-/-^ mice were purchased from Shulaibao (Wuhan) Biotechnology Co., Ltd, and were administrated with β-aminopropionitrile (BAPN, 1g/L, Sigma-Aldrich, St Louis) in the drinking water for 3 weeks followed by 2500 ng/kg/min Angiotensin II (AngII, Sigma-Aldrich, St Louis) subcutaneous pump for 2 weeks to induce AD [27]. Selective integrin β3 inhibitors Cyclo(-RGDfK) (MedChemExpress, Houston, TX, USA) were administered synchronously with 2500 ng/kg/min AngII subcutaneous pump at dosage of 5 mg/kg/d by intraperitoneal injection, while Cyclo(-RADfK) (MedChemExpress, Houston, TX, USA) was injected as negative control. The mice were euthanized, following which the aortic tissues were harvested and subsequently properly stored.

### Histology, immunohistochemistry and immunofluorescence

All mouse aortas harvested from animal experiments were fixed in 4% paraformaldehyde for 48 hours, followed by embedded in paraffin, and then sliced into 4 μm sections. Aortic morphology was assessed by hematoxylin and eosin (H&E) staining and the integrity of aortic wall elastin was evaluated by van Gieson (EVG) staining. Images were captured using an OLYMPUS BX53

Microscope (OLYMPUS, Beijing, China). Immunohistochemistry was observed using an OLYMPUS BX53 Microscope (OLYMPUS, Beijing, China) after ICAM-1 (1:100, Proteintech, 10831-1-AP), VCAM-1 (1:100, Proteintech, 83719-1-RR), NOX2 (1:100, Abclonal, A19701), NOXO2 (1:100, Abclonal, A1178), NOXA2 (1:100, Abclonal, A1148), cleaved caspase3 (1:100, Abclonal, A27145) and β-catenin (1:100, Abclonal, A11512) staining. The ImageJ software was utilized for the analysis of Optical Density (OD) values of the images [28]. Immunofluorescence was observed by OLYMPUS BX51 Microscope (OLYMPUS, Beijing, China) after AP5 (1:100, kerafast, EBW107) / CD31 (1:100, Servicebio, GB120005-100) staining and then 4’, 6-diamidinao-2-phenylindole (DAPI, Beyotime, C1002) was used to counterstain the nucleus [29].

### Cell culture

HUVEC (CRL-1730) and Human Embryonic Kidney 293T (293T, CRL-3216) were purchased from the American Type Culture Collection (ATCC; Manassas, VA, USA). HUVECs were cultured in Roswell Park Memorial Institute (RPMI) 1640 medium supplemented with 10% fetal bovine serum (FBS; Hyclone Laboratories, Inc., Logan, UT, USA) and 1% penicillin/streptomycin (NCM Biotech), and 293T cells were cultured in Dulbecco’s modified Eagle’s medium (DMEM) containing 4.5 g/L glucose supplemented with 10% FBS and 1% penicillin/streptomycin. All cell lines were cultured in a humidified 37°C incubator with 95% air and 5% CO_2_ [30].

### Cellular immunofluorescence

HUVEC cells were inoculated into confocal dish (In Vitro Scientific, D35-14-1-N) and treated with AngII and Cyclo(-RADfK). Cells were fixed with 4% paraformaldehyde for 20 min and permeabilized with 0.3% TritonX-100 for 15 min at room temperature. After blocking with 5% BSA, cells were incubated with primary antibody against AP5 (1:100, kerafast, EBW107), integrin β3 (1:100, Abclonal, A24844) and AT1R (1:100, Abclonal, A4140) at 4°C overnight, then the FITC-conjugated and CY3-conjugated secondary antibody and DAPI were used for visualization. Fluorescent images were captured by OLYMPUS FV1000 Microscope (OLYMPUS, Beijing, China).

### Immunoprecipitation

293T cells were transfected with plasmids for 24 hours, and lysed with cell lysis buffer for Western and IP (P0013; Beyotime, Shanghai, China). Lysates were preincubated with 20 μl protein A/G agarose beads (Santa Cruz, sc-2003) for 30 min to eliminate nonspecific binding and incubated with labelling antibody against GFP (Abclonal, AE078) for 1 h at 4 °C. The antigen-antibody complex was precipitated by protein A/G agarose beads and collected for immunoblotting to check the interaction between integrin β3 with AT1R, integrin β3 with GSK3β.

### Western Blot

HUVEC cells were harvested after various treatments. Subsequently, the cells were collected and washed with cold PBS for three times before being lysed in RIPA buffer supplemented with protease and phosphatase inhibitor cocktails (HY-K0010, HYK0021; MedChemExpress, Houston, TX, USA) to obtain whole cell lysates. Lysates were used for electrophoresis and transferred onto 0.45 μm polyvinylidene fluoride (PVDF) membrane. The membranes were blocked with 5% BSA and incubated with specific primary antibodies overnight at 4°C. HRP-conjugated secondary antibodies against corresponding species (1:2000, Abclonal, AS014 (anti-rabbit) and AS003 (anti-mouse)) and enhanced chemiluminescence kit (Boster, AR1171) were used for detection.

The following primary antibodies were used for immunoblotting at 1:1000 dilution: FAK (Cell Signaling Technology, 3285), p-FAK (Tyr397) (Santa Cruz, sc-81493), ICAM-1 (Proteintech, 10831-1-AP), VCAM-1 (Proteintech, 83719-1-RR), NOX2 (Abclonal, A19701), NOXO2 (Abclonal, A1178), NOXA2 (Abclonal, A1148), cleaved caspase3 (Abclonal, A27145), integrin β3 (Abclonal, A24844), AT1R (Abclonal, A4140), β-catenin (Abclonal, A11512), GSK3β (Abclonal, A2081), p-GSK3β (S9) (Abclonal, AP1088) and GAPDH (Proteintech, 60004-1-lg).

### Dihydroethidium staining

HUVEC were washed with cold PBS and subsquently incubated with 1 μM Dihydroethidium (DHE, S0063, Beyotime, Shanghai, China) and Hoechst (Beyotime, C1027) for 30 min at 37 °C. The fluorescence was measured utilizing a Fluorescence microscope (OLYMPUS, Beijing, China). And the mean fluorescence intensity was quantitative analyzed by Image J software.

### TUNEL staining assay

After various interventions, HUVEC were fixed in 4% paraformaldehyde for 20 minutes at room temperature and permeabilized with 0.3% TritonX-100 at 37°C for 15 minutes. Appropriate volume of TUNEL reaction solution (C1086, Beyotime, Shanghai, China) was applied and the reaction was kept at 37°C for 1 hour. The fluorescence was measured as previously mentioned.

### F-actin staining assay

HUVEC were fixed in 4% paraformaldehyde for 10 minutes at room temperature and permeabilized with 0.1% TritonX-100 at 37°C for 20 minutes. Appropriate volume of Actin-Tracker Red (1:100, C2207S, Beyotime, Shanghai, China) was applied and the reaction was kept at 37°C for 1 hour. The fluorescence was measured as previously mentioned.

### Statistical analysis

Data are presented as mean ± standard deviation (SD). Statistical analysis was conducted with Graphpad Prism software (version 10.0.0). For quantitative data, Shapiro-Wilk test was first performed to test normal distribution and F test was used to compare variances. Unpaired *t*-test was used to analyze the differences between two groups and one-way analysis of variance (ANOVA) combined with Tukey multiple comparisons was used when there were more than two groups. Nonparametric test (Mann-Whitney test or Kruskal-Wallis test) was used for data not conforming to normal distribution. For qualitative data and rank data, Fisher’s exact test was used. Statistical significance was defined as *p*<0.05.

## Result

### Integrin β3 Activation is Elevated in Experimental Aortic Dissection

To determine the role of integrin β3 in the pathogenesis of aortic dissection (AD), we analyzed the publicly available single-cell RNA sequencing (scRNA-seq) dataset (GSE222318) containing tissue samples from different stages of AD. Our analysis revealed a significant upregulation of integrin β3 activation, particularly in endothelial cells (ECs), during the acute phase of AD (Supplementary Figures 1A-B). Given that integrins lack intrinsic kinase activity and depend on downstream kinases for activation, we examined the expression of focal adhesion kinase (FAK) and proline-rich tyrosine kinase 2 (Pyk2), two major tyrosine kinases associated with integrin signaling [31–33]. scRNA-seq data showed a substantial increase in the activation of these kinases (Figures 1A-B, Supplementary Figure 1C).

**Figure 1.**
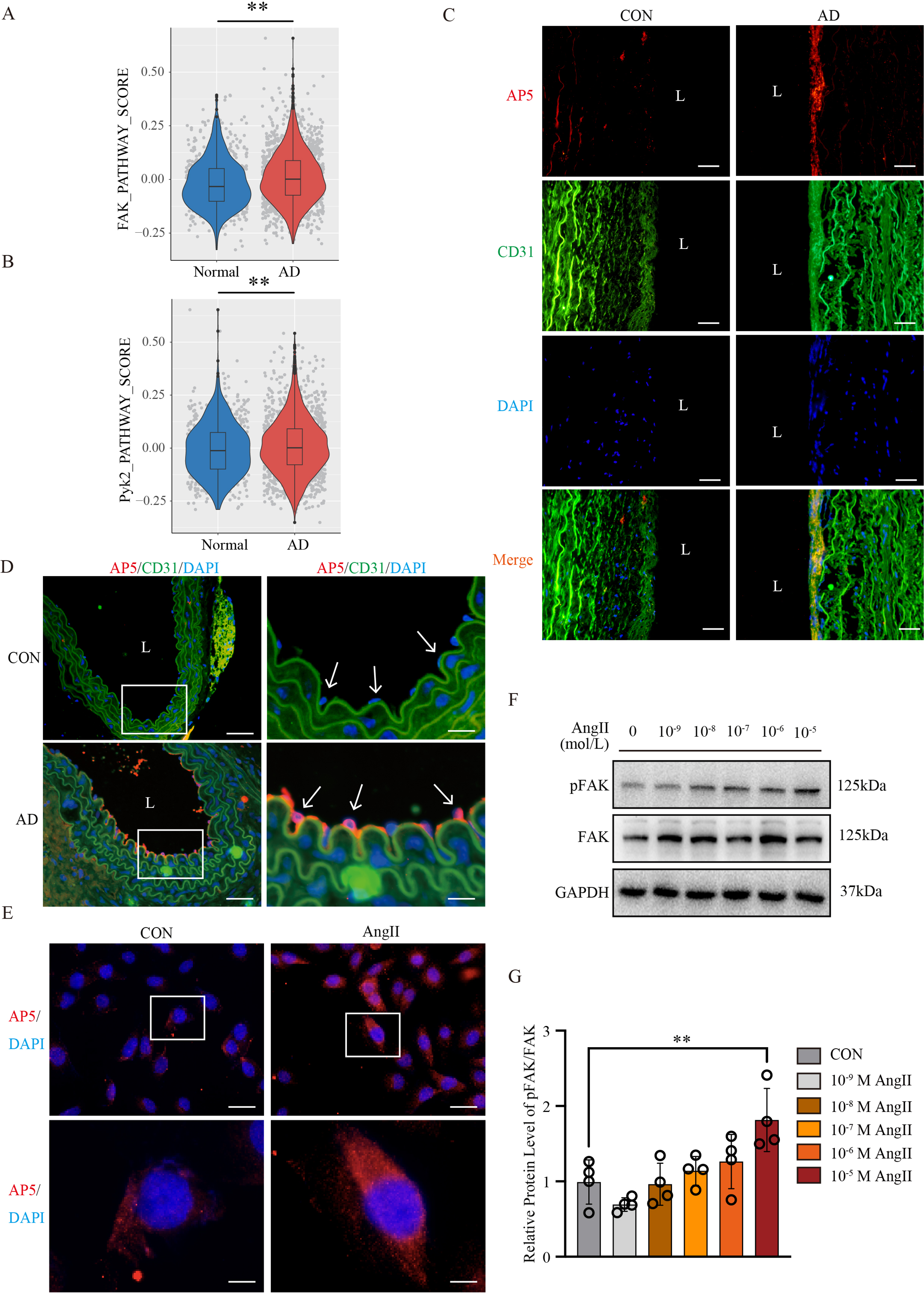
Integrin β3 Activation is Elevated in Experimental Aortic Dissection. (A-B) FAK pathway and PYK2 pathway scores in the Normal and AD groups. (C) Exemplary images of AP5/CD31 IF staining of human aortic tissue. (D) Exemplary images of AP5/CD31 IF staining of mouse aortic tissue. (E) Immunofluorescence staining showing enhanced activation of Integrin β3 in Ang II-stimulated HUVEC cells. (F-G) Representative immunoblots showing the protein abundance of pFAK and FAK in HUVEC cells treated with AngII, GAPDH served as loading control (F), and bar graph summarizing the results of pFAK/FAK (G). At least three independent experiments were performed. All data were represented as the mean ± SD, ** *p* < 0.01, One-way ANOVA.

To validate these findings in human AD, we performed immunofluorescent staining of human aortic specimens using the active conformation-specific integrin β3 antibody (AP5) [34]. Our results demonstrated significantly increased AP5 fluorescence in aortic ECs, suggesting heightened integrin β3 activation in AD (Figure 1C). To further investigate this phenomenon, we induced AD in ApoE^-/-^ mice by administering β-aminopropionitrile (BAPN) followed by angiotensin II (AngII) infusion (Supplementary Figure 2), as described previously [27]. Immunofluorescence analysis of these mice showed a marked increase in AP5 staining in the aortic endothelium, corroborating our human tissue findings (Figure 1D).

Furthermore, *in vitro* experiments using human umbilical vein endothelial cells (HUVECs) exposed to AngII showed a dose-dependent increase in AP5 expression, indicating AngII-mediated integrin β3 activation. This was accompanied by increased phosphorylation of FAK, suggesting crosstalk between AT1R and integrin β3 signaling (Figures 1E-G). Taken together, these findings establish integrin β3 as a key player in AD progression and suggest that its inhibition may represent a novel therapeutic target.

### Pharmacological Inhibition of Integrin β3 Attenuates AD Progression

To specifically inhibit integrin β3 activation in AD models, we used a specific cyclic peptide Cyclo(-RGDfK). Male 6-week-old ApoE^-/-^ mice were randomly allocated into 3 groups, namely the control group, AngII combined with 5 mg/kg/d Cyclo(-RADfK) group, AngII combined with 5 mg/kg/d Cyclo(-RGDfK) group (Figure 2A). Notably, Cyclo(-RGDfK) treatment significantly improved survival rates compared to Cyclo(-RADfK) -treated group (Figures 2B, 2D). Additionally, histological analyses revealed a substantial reduction in AD incidence and lesion severity in the Cyclo(-RGDfK)-treated group (Figures 2C, 2E). Elastin fragmentation and extracellular matrix (ECM) degradation are hallmark features of AD. Van Gieson (EVG) staining demonstrated that Cyclo(-RGDfK) treatment preserved elastin integrity and reduced aortic dilation compared to Cyclo(-RADfK) -treated mice (Figures 2F-H). These results indicate that integrin β3 inhibition mitigates AD progression by preserving aortic wall structure and preventing dissection.

**Figure 2.**
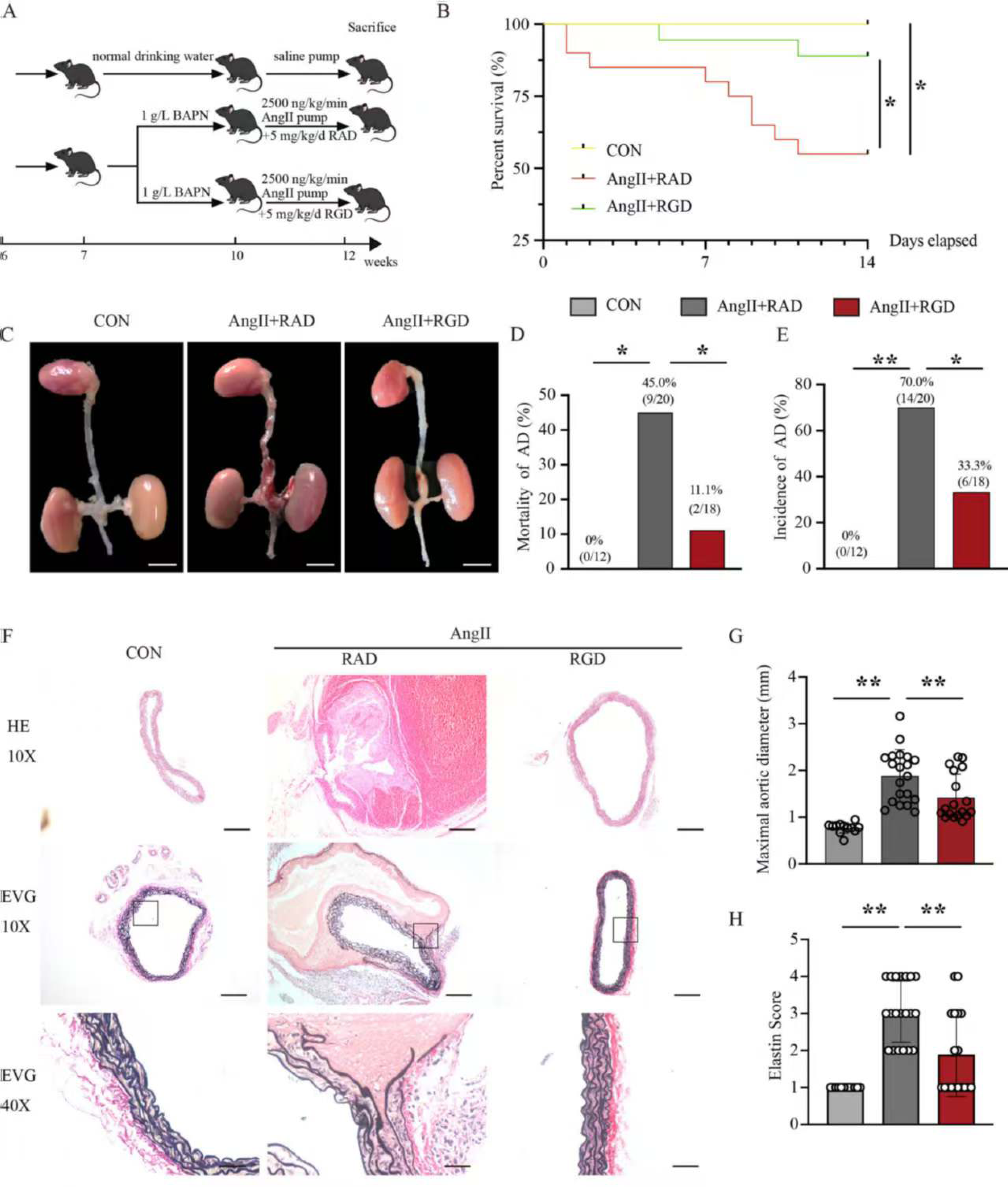
Pharmacological Inhibition of Integrin β3 Attenuates AD Progression. (A) Schematic diagram of animal experiment procedure. (B) Kaplan-Meier survival curve of AngII-induced AD model mice with or without Cyclo(-RGDfK) treatment. (C) Representative macroscopic images of mice aorta isolated from the indicated groups. (D) Mortality rate of AngII-administrated mice with or without Cyclo(-RGDfK) treatment. Fisher’s exact test was used for statistical analysis. (E) Incidence of AD in AngII-treated mice with or without Cyclo(-RGDfK) treatment. Fisher’s exact test was used for statistical analysis. (F) Exemplary images of HE staining and EVG staining of the aortic tissues from the above mentioned groups. (G) Quantification of maximum aortic diameter. Brown-Forsythe ANOVA test was used for statistical analysis. (H) Elastin score assessed by EVG staining to evaluate elastin degradation in AngII-induced and Cyclo(-RGDfK) treated mouse aortas, Fisher’s exact test was used for statistical analysis. N = 12/20/18, * *p* < 0.05, ** *p* < 0.01.

### Integrin β3 Drives Endothelial Dysfunction via ROS and Apoptosis

Integrin signaling is known to regulate cytoskeletal remodeling, apoptosis, and reactive oxygen species (ROS) production [27, 35, 36]. To assess whether integrin β3 inhibition protects ECs, we analyzed inflammatory markers in the aorta of AD mice. Cyclo(-RGDfK) treatment significantly reduced endothelial activation markers ICAM-1 and VCAM-1, as well as oxidative stress indicators NOX2, NOXO2, and NOXA2. Consistently, Cyclo(-RGDfK) reduced apoptosis, as indicated by decreased cleaved caspase-3 expression (Figure 3A, Supplementary Figure 3A). *In vitro*, AngII-treated HUVECs exhibited elevated ROS production and apoptosis, which were significantly alleviated by Cyclo(-RGDfK) treatment (Figures 3B-I). Furthermore, F-actin staining revealed that AngII disrupted endothelial cytoskeletal integrity, whereas Cyclo(-RGDfK) preserved stress fiber organization (Figure 3J). These data indicate that integrin β3 inhibition effectively prevents AngII-induced endothelial dysfunction, a key contributor to AD development [37].

**Figure 3.**
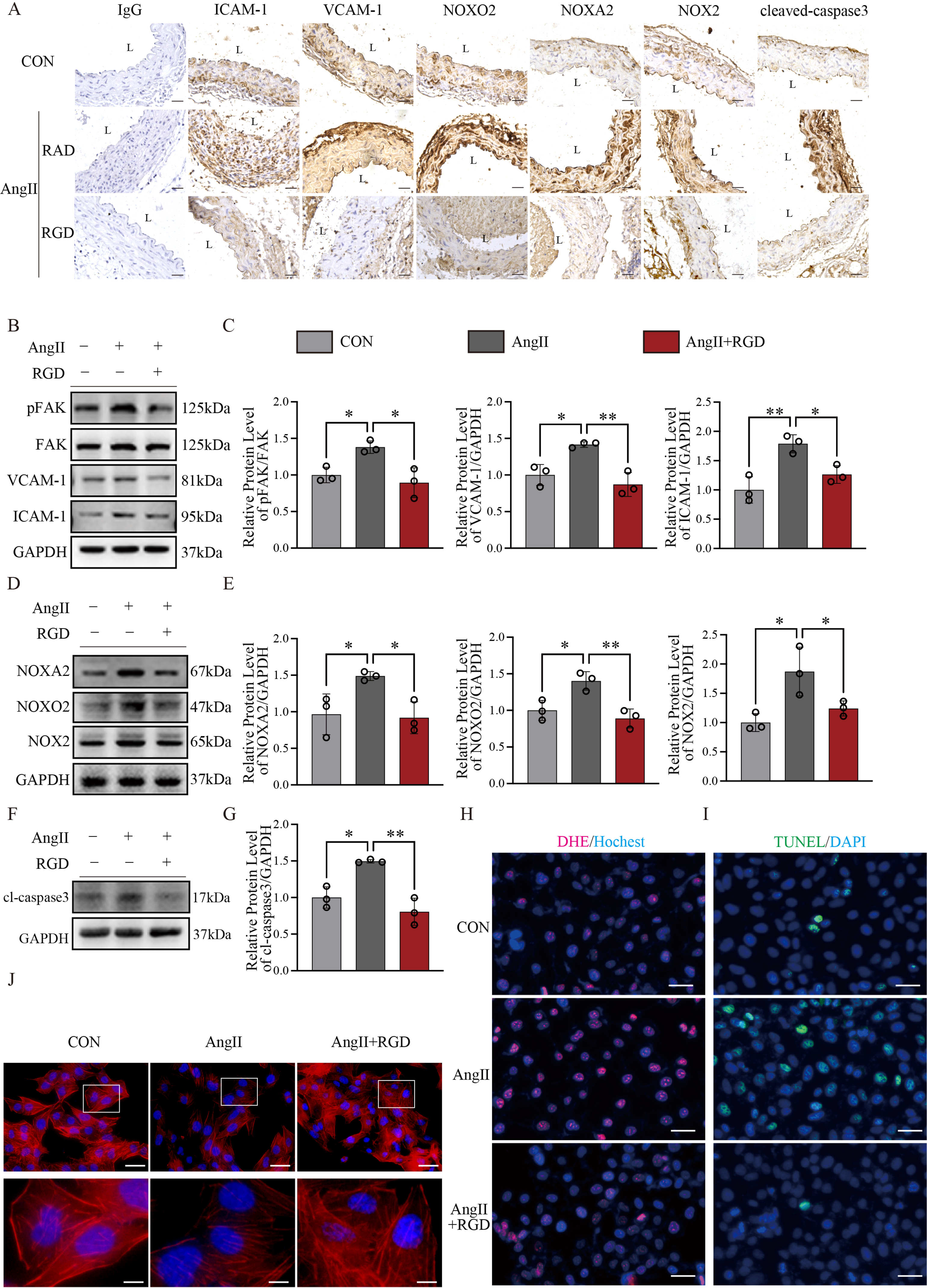
Integrin β3 Drives Endothelial Dysfunction via ROS and Apoptosis. (A) Representative ICAM-1, VCAM-1, NOX2, NOXO2, NOXA2 and cleaved-caspase3 IHC staining of mice aortic tissue from the indicated groups. Non-specific IgG served as negative control. (B-C) Representative immunoblots showing the protein abundance of pFAK, FAK, ICAM-1 and VCAM-1 in HUVEC cells treated with AngII in combination with Cyclo(-RGDfK), GAPDH served as loading control (B), and bar graph summarizing the results (C).(D-E) Representative immunoblots showing the protein abundance of NOX2, NOXO2 and NOXA2 in HUVEC cells treated with AngII in combination with Cyclo(-RGDfK), GAPDH served as loading control (D), and bar graph summarizing the results (E). (F-G) Representative immunoblots showing the protein abundance of cleaved-caspase3 in HUVEC cells treated with AngII in combination with Cyclo(-RGDfK), GAPDH served as loading control (F), and bar graph summarizing the results (G). (H-J) Typical images of DHE staining (H), TUNEL staining (I) and F-actin staining (J) of HUVEC cells after the intervention with AngII and Cyclo(-RGDfK). At least three independent experiments were performed. All data were represented as the mean ± SD, * *p* < 0.05, ** *p* < 0.01, One-way ANOVA.

### AT1R-Integrin β3 Crosstalk Amplifies Endothelial Injury

To explore the molecular crosstalk between AT1R and integrin β3, we performed co-immunoprecipitation (Co-IP) assays, revealing a direct interaction between integrin β3 and AT1R in endothelial cells. Notably, AngII stimulation enhanced this interaction, suggesting that AT1R activation facilitates integrin β3 recruitment (Figures 4A-C). Besides, Immunofluorescence further confirmed co-localization of integrin β3 and AT1R in HUVECs (Figure 4D).

**Figure 4.**
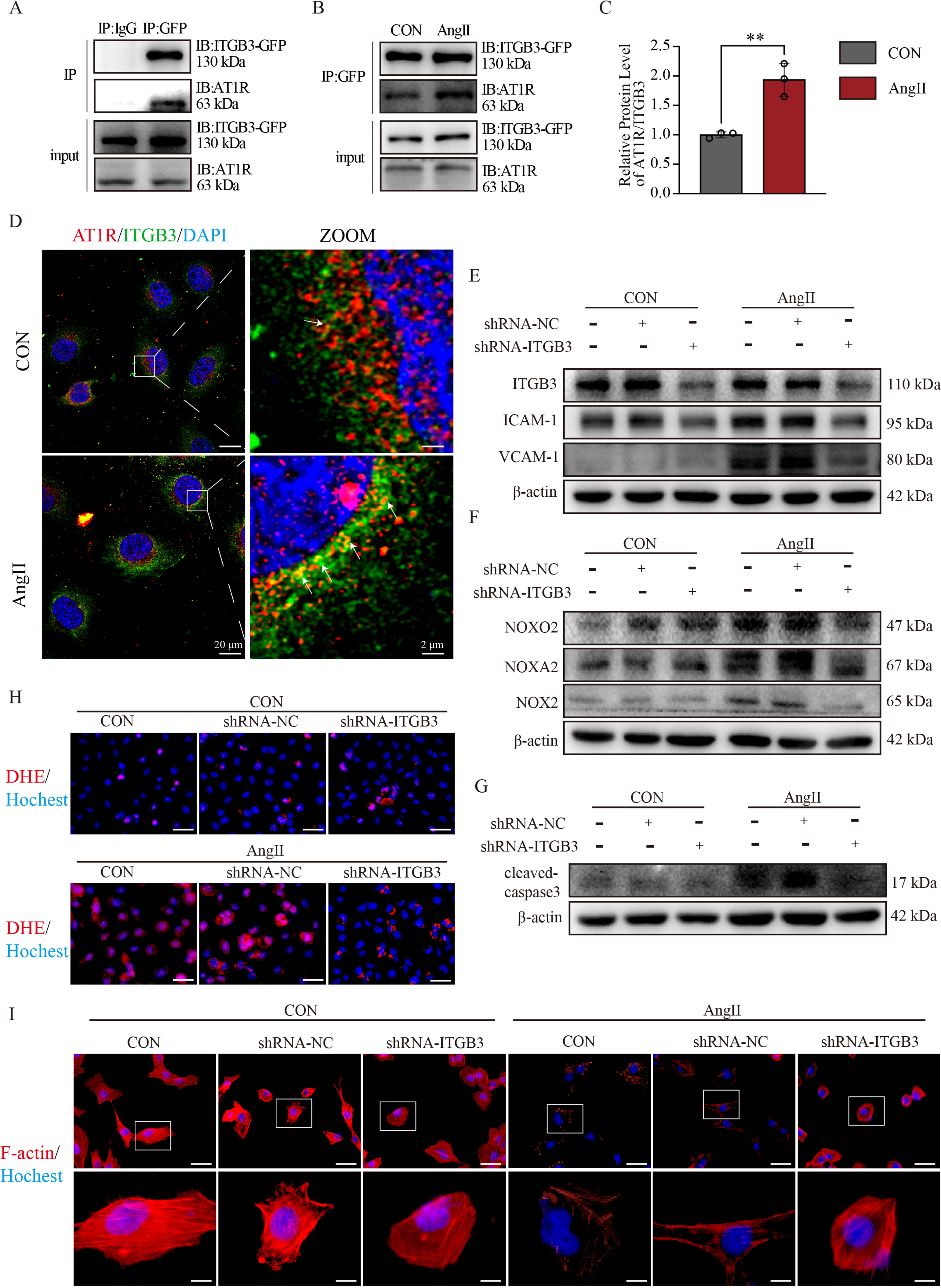
AT1R-Integrin β3 Crosstalk Amplifies Endothelial Injury. (A) Representative Co-IP showing the interaction between Integrin β3 and AT1R (B-C) Representative Co-IP and quantification showing the effect of Ang II stimulation on the interaction between Integrin β3 and AT1R. (D) Exemplary images of Integrin β3 and AT1R IF staining of HUVEC cells with and without AngII stimulation. (E-G) HUVEC cells were infected with lentivirus containing Integrin β3 shRNA or negative control, and then stimulated with or without Ang II at 10^-5^ mol/L for 24 h. Western blot analysis were performed to assess the protein level of ICAM-1, VCAM-1, NOX2, NOXO2, NOXA2 and cleaved-caspase3. (H-I) Typical images of DHE staining (H) and F-actin staining (I) of HUVEC cells after the infection with lentivirus containing Integrin β3 shRNA or negative control, and then stimulated with or without Ang II. At least three independent experiments were performed. All data were represented as the mean ± SD, ** *p* < 0.01, *t*-test.

To determine whether integrin β3 activation mediates endothelial injury, we knocked down integrin β3 in HUVECs and observed that AngII-induced upregulation of ICAM-1, VCAM-1, and ROS markers was significantly attenuated (Figures 4E-G, Supplementary Figure 4A). DHE staining, TUNEL staining and F-actin staining further confirmed reduced oxidative stress, apoptosis and damage of cytoskeleton in integrin β3-deficient cells (Figures 4H-I, Supplementary Figure 4B).

To confirm the functional relevance of integrin β3 activation, we treated ECs with manganese chloride (MnCl_2_), which enhances integrin activation [38, 39]. MnCl_2_ treatment exacerbated ROS accumulation and cytoskeletal alterations in AngII-treated cells (Supplementary Figure 5A). Conversely, co-treatment with losartan (AT1R inhibitor) and Cyclo(-RGDfK) robustly suppressed integrin β3 activation and associated endothelial injury (Supplementary Figures 5B-E). Together, these results establish integrin β3-AT1R crosstalk as a critical mediator of AngII-induced endothelial dysfunction.

### Non-Canonical Wnt/β-Catenin Signaling Mediates Integrin β3-Driven EC Injury

To delineate downstream pathways involved in integrin β3-mediated endothelial injury, we performed RNA sequencing (RNA-Seq) analysis in integrin β3 knockdown ECs. KEGG pathway enrichment analysis and gene set enrichment analysis (GSEA) identified significant alterations in the Wnt signaling pathway, implicating Wnt dysregulation in integrin β3-driven endothelial dysfunction (Figures 5A-C) [40, 41].

**Figure 5.**
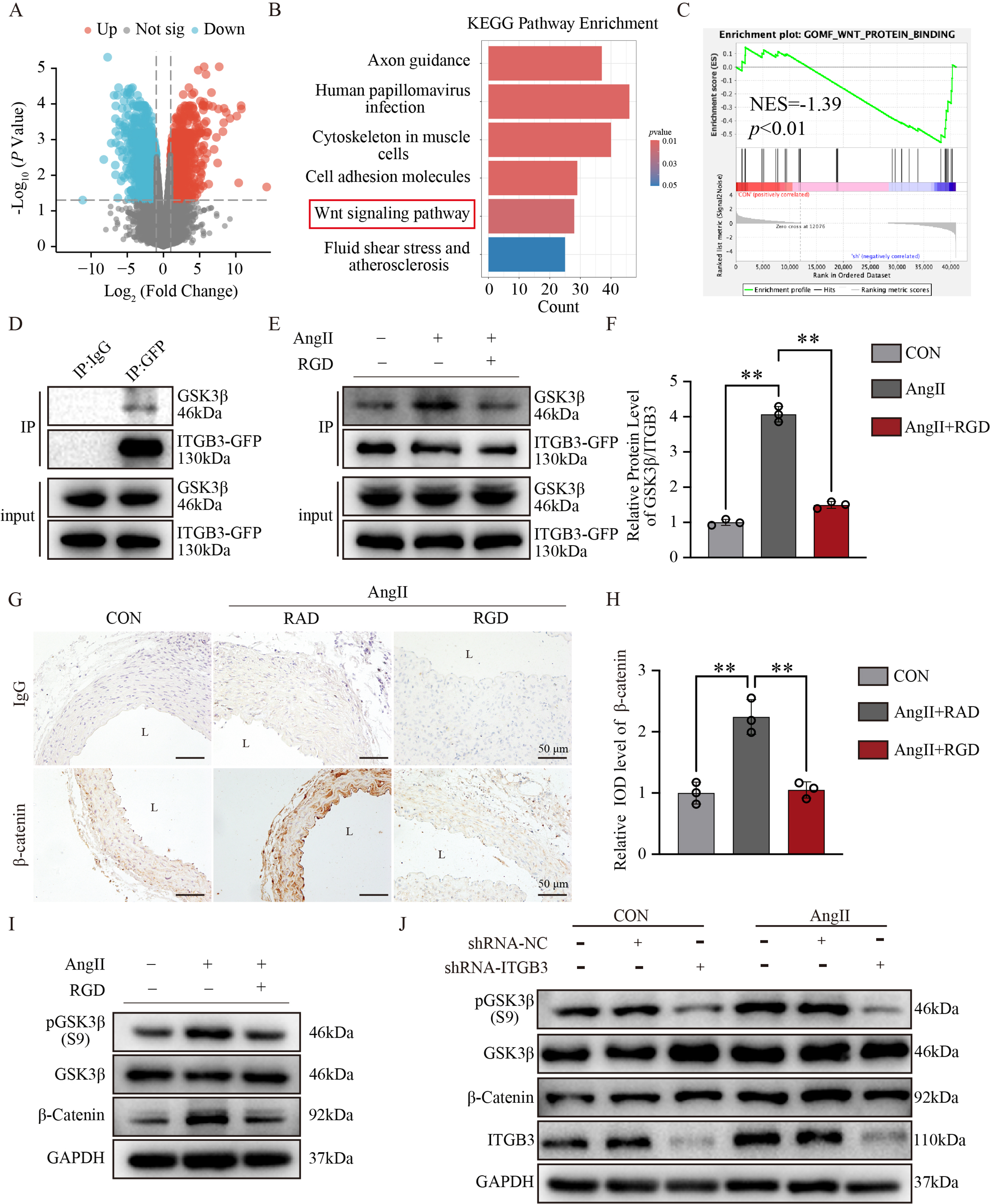
Non-Canonical Wnt/β-Catenin Signaling Mediates Integrin β3-Driven EC Injury. (A-C) RNA sequencing analysis after integrin β3 knockdown in HUVEC cells. DEGs analysis (A) and KEGG pathway enrichment analysis (B) showed that Wnt signaling pathway was significantly changed after integrin β3 was knockdown, and GSEA (C) also confirmed that Wnt signaling pathway was significantly changed after integrin was β3 knockdown. (D) Representative Co-IP showing the interaction between Integrin β3 and GSK3β (E-F) Representative Co-IP and quantification showing the effect of Ang II stimulation on the interaction between Integrin β3 and GSK3β. (G-H) Representative β-catenin IHC staining of mice aortic tissue from the indicated groups. Non-specific IgG served as negative control. The staining intensity of GSK3β was summarized in the bar graph. (I) Representative immunoblots showing the protein abundance of GSK3β, p-GSK3β and β-catenin in HUVEC cells treated with AngII in combination with Cyclo(-RGDfK), GAPDH served as loading control. (J) HUVEC cells were infected with lentivirus containing Integrin β3 shRNA or negative control, and then stimulated with or without Ang II. Western blot analysis were performed to assess the protein level of GSK3β, p-GSK3β and β-catenin. At least three independent experiments were performed. All data were represented as the mean ± SD, * *p* < 0.05, ** *p* < 0.01, One-way ANOVA.

Canonical Wnt signaling typically involves Frizzled-LRP5/6 receptor activation, leading to β-catenin stabilization [42, 43]. However, pharmacological inhibition of canonical Wnt signaling (LGK974) failed to suppress β-catenin accumulation in AngII-treated ECs (Supplementary Figures 6A-C). Additionally, AKT inhibition, another alternative Wnt regulatory pathway [44, 45], did not attenuate β-catenin upregulation (Supplementary Figures 6D-F).

Intriguingly, Co-IP studies revealed a direct interaction between integrin β3 and glycogen synthase kinase 3β (GSK3β), which was further enhanced by AngII treatment (Figures 5D-F). Immunohistochemistry confirmed elevated β-catenin expression in AD mouse aortas, which was significantly reduced by Cyclo(-RGDfK) (Figures 5G-H). Western blot analysis in ECs demonstrated that Cyclo(-RGDfK) or integrin β3 knockdown markedly suppressed GSK3β phosphorylation and β-catenin accumulation (Figures 5I-J, Supplementary Figures 7A-B). These findings establish a novel signaling axis wherein integrin β3 directly interacts with GSK3β to activate non-canonical Wnt signaling, contributing to AD pathogenesis.

Taken together, our study provides compelling evidence that integrin β3-AT1R crosstalk is a key mediator of AD progression. We uncover a novel molecular mechanism wherein AT1R activation triggers integrin β3 activation, leading to endothelial injury *via* non-canonical Wnt signaling. Dual inhibition of AT1R and integrin β3 may be a promising therapeutic strategy.

## Discussion

AD is a catastrophic cardiovascular event characterized by the progressive degeneration of the aortic wall, leading to intimal tearing and fatal complications. Despite advancements in surgical and pharmacological interventions, effective medical therapies for AD remain elusive. In this study, we identify integrin β3 as a novel pathogenic driver of AD and elucidate a previously unrecognized crosstalk between AT1R and integrin β3 in mediating endothelial dysfunction. Mechanistically, we demonstrate that AT1R activation enhances integrin β3 signaling, which in turn triggers non-canonical Wnt pathway activation *via* GSK3β interaction, ultimately leading to endothelial cell injury and aortic wall degeneration (Figure 6). These findings highlight a novel therapeutic target for AD and advocate for a dual-target approach involving both AT1R and integrin β3 inhibition.

**Figure 6.**
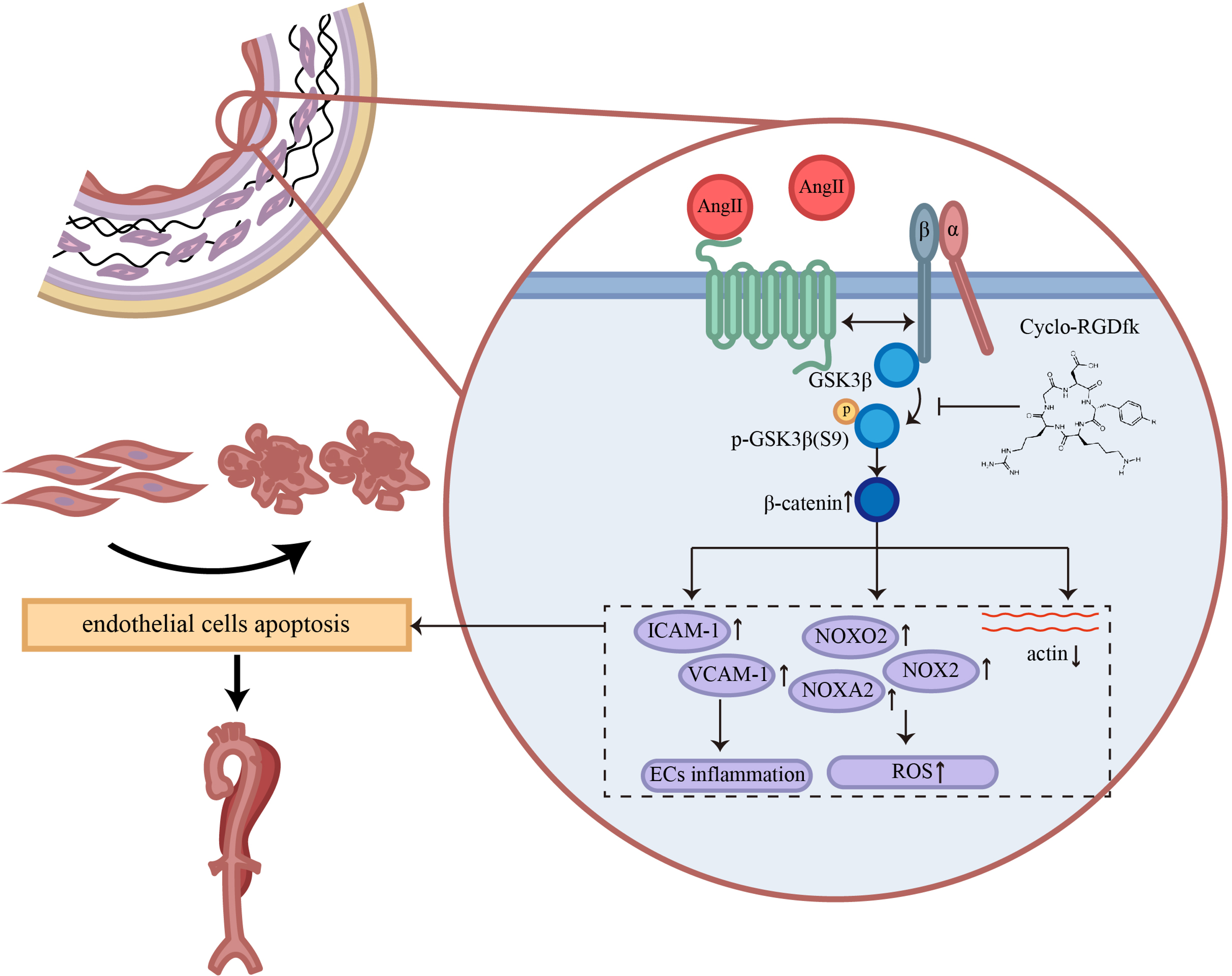
Graphical Abstract. | AngII stimulation orchestrates the activation of AT1R, which subsequently facilitates its interplay with integrin β3, culminating in integrin β3 activation during AD pathogenesis. Furthermore, activated integrin β3 engages with GSK3β, triggering the activation of the non-canonical Wnt signaling pathway, which subsequently upregulates the expression of β-catenin, ultimately driving vascular endothelial cell injury and promoting AD occurrence.

### Integrin β3 as a Key Pathogenic Mediator in AD

Emerging evidence suggests that integrins, particularly integrin β3, play critical roles in vascular homeostasis and disease. Integrin β3 is a well-characterized transmembrane receptor that regulates cell adhesion, migration, and survival [9, 10]. In the cardiovascular system, integrin β3 modulates endothelial barrier function and vascular remodeling [11, 12]. Previous studies have linked integrin β3 to aortic aneurysm formation [46], yet its role in AD remained undefined. Our study demonstrates that integrin β3 is significantly upregulated in ECs during AD progression, with its activation closely associated with increased FAK and Pyk2 signaling. Importantly, pharmacological inhibition of integrin β3 using Cyclo(-RGDfK) effectively attenuated AD pathogenesis, reduced endothelial apoptosis, and preserved aortic wall integrity.

These findings expand on previous work showing that integrin β3 signaling is critical for vascular endothelial function [13]. A recent study highlighted the role of integrin β3 in regulating MMPs, which contribute to ECM degradation and vascular instability [14, 15]. Our study builds on this evidence by demonstrating that integrin β3 activation exacerbates endothelial dysfunction and promotes AD, potentially through a novel interaction with AT1R.

### Crosstalk Between AT1R and Integrin β3 in AD Progression

The renin-angiotensin system (RAS) is a well-established regulator of vascular pathology, with AT1R activation playing a pivotal role in AD pathogenesis [16, 17]. Despite this, clinical trials using ACEis and ARBs have shown only modest benefits in reducing AD risk [18]. This suggests that AT1R signaling alone may not fully account for the complex molecular mechanisms underlying AD progression.

Our study identifies integrin β3 as a critical mediator of AT1R-driven endothelial injury. We show that AT1R activation enhances integrin β3 expression and promotes its interaction with AT1R, thereby amplifying endothelial dysfunction. This is supported by our co-immunoprecipitation and immunofluorescence studies, which confirm a direct interaction between these two receptors. Furthermore, pharmacological inhibition of either AT1R or integrin β3 partially attenuates endothelial injury, while combined inhibition provides greater protective effects. These findings suggest that integrin β3 functions as an essential downstream effector of AT1R, contributing to AD pathogenesis through a synergistic signaling axis.

Previous studies have demonstrated that integrins interact with various GPCRs, including VEGFR and EGFR [19–21]. Our findings extend this knowledge by demonstrating a direct interplay between integrin β3 and AT1R, offering a novel mechanistic insight into how AT1R signaling contributes to AD. Given the limited efficacy of AT1R blockade alone, our findings highlight the potential for dual-target therapy as a more effective strategy in AD treatment.

### Non-Canonical Wnt Signaling as a Novel Pathogenic Pathway in AD

The Wnt signaling pathway plays a fundamental role in vascular development and endothelial homeostasis [40, 41]. While the canonical Wnt pathway involves β-catenin activation through Frizzled-LRP5/6 receptors, non-canonical Wnt signaling operates independently of these receptors and is less well understood [42, 47]. In our study, RNA-Seq analysis identified significant enrichment of Wnt pathway genes following integrin β3 activation. Surprisingly, inhibition of canonical Wnt signaling with LGK974 failed to prevent β-catenin accumulation, suggesting the involvement of a non-classical pathway.

Our co-immunoprecipitation experiments revealed a direct interaction between integrin β3 and GSK3β, which was further enhanced by AngII stimulation. This interaction resulted in increased GSK3β phosphorylation and β-catenin stabilization, independent of Frizzled receptor activation. These findings unveil a novel integrin β3-GSK3β axis that regulates endothelial dysfunction in AD.

Prior studies have implicated non-canonical Wnt signaling in various vascular diseases, including atherosclerosis and pulmonary hypertension [47, 48]. However, its role in AD has remained unexplored. Our findings provide the first evidence that integrin β3-mediated non-canonical Wnt signaling contributes to endothelial dysfunction and AD pathogenesis. Given that β-catenin plays a central role in vascular homeostasis, its dysregulation may represent a key driver of AD progression.

### Therapeutic Implications and Future Directions

Our study provides compelling evidence that integrin β3 is a promising therapeutic target for AD. The current standard of care for AD remains surgical repair and antihypertensive therapy, with limited pharmacological options available. The discovery of integrin β3 as a critical mediator of AD opens new avenues for therapeutic intervention.

Our findings suggest that dual inhibition of AT1R and integrin β3 may offer superior efficacy in AD management. Given that integrin β3 is a mechanosensitive receptor, future studies should investigate its role in flow-dependent vascular remodeling. Additionally, further research is needed to determine whether integrin β3 inhibitors can be safely translated into clinical practice.

While our study provides novel insights into AD pathogenesis, several limitations warrant consideration. First, our experiments were conducted primarily in AngII-induced AD models, which may not fully capture the heterogeneity of human AD. Second, the structural details of the AT1R-integrin β3 and integrin β3-GSK3β interactions remain unknown. High-resolution structural studies, such as cryo-EM or AlphaFold2-based modeling [49], could provide deeper mechanistic insights. Finally, long-term studies are needed to assess the safety and efficacy of integrin β3-targeted therapies *in vivo*.

## Conclusion

In conclusion, our study identifies integrin β3 as a novel driver of AD and elucidates a previously unrecognized AT1R-integrin β3-GSK3β signaling axis. We demonstrate that integrin β3 activation exacerbates endothelial dysfunction through non-canonical Wnt signaling, contributing to AD pathogenesis. Pharmacological inhibition of integrin β3 attenuates AD progression, highlighting a promising new therapeutic strategy. These findings provide a strong rationale for dual targeting of both AT1R and integrin β3 in AD and lay the foundation for future translational research aimed at developing more effective treatments for this life-threatening disease.

## Declarations

### Ethics approval and consent to participate

The authors declare the ethics approval has been acquired and this manuscript has been approved by all authors for publication.

## Availability of data and material

The original contributions presented in the study are included in the article and the Supplementary Material, further inquiries can be directed to the corresponding authors.

## Authors’ contribution

Hesong Zeng: Conceptualization, Methodology, Funding acquisition Hongjie Wang: Supervision, Project administration

Hongcheng Jiang and Zixuan Liu: Software, Validation, Formal analysis, Data Curation, Investigation, Visualization, Writing - Original Draft

Yu Li and Xiaodan Zhong: Investigation, Formal analysis, Data Curation, Writing - Review & Editing

Yunkun Qu and Lei Dai: Data Curation

Ying zhang, Mengwen Wang, Wenjun Zhang, Wanjun Liu and Xingwei He: Investigation Shiliang Li: Resources

Zhiyong Li, Wei Dong and Thati Madhusudhan: Writing - Review & Editing

## Acknowledgements

We thank Experimental Medicine Center of Tongji Hospital, Tongji Medical School, Huazhong University of Science and Technology for providing OLYMPUS BX53 Microscope.

## Non-standard Abbreviations and Acronyms

ACEi: Angiotensin Converting Enzyme inhibitor
AD: Aortic Dissection
AngII: Angiotensin II
ARB: Angiotensin Receptor Blocker
AT1R: angiotensin II type 1 receptor
ARRIVE: Animal Research: Reporting of In Vivo Experiments
ANOVA: analysis of variance
BAPN: β-aminopropionitrile monofumarate
BSA: Bovine Serum Albumin
DAPI: 4’, 6-diamidinao-2-phenylindole
DMSO: Dimethyl Sulfoxide
DEGs: Differentially Expressed Genes
EC: Endothelial Cell
EVG: Evaluated by Van Gieson
ECM: Extracellular Matrix
FDR: False discovery rate
FBS: Fetal Bovine Serum
GPCR: G Protein-Coupled Receptor
GSEA: Gene Set Enrichment Analysis
GSK3β: Glycogen Synthase Kinase 3β
HUVEC: Human Umbilical Vein Endothelial Cell
MMP9: Matrix Metalloproteinase-9
NES: Normalized Enrichment Score
NOX: NADPH Oxidase
OD: Optical Density
PBS: Phosphate-Buffered Saline
PAR: Protease-Activated Receptors
ROS: Reactive Oxidative Stress
SD: Standard Deviation

